# Single-Cell full-length Isoform Sequencing Unveils Transcriptional Dysregulation in Autism Spectrum Disorder During Cerebral Cortex Development

**DOI:** 10.1101/2024.12.04.626807

**Authors:** Xiaoyi Xu, Jun Wang, Kunhua Hu, Dan Su, Qingpei Huang, Xiaotang Fan, Xiaoying Fan

## Abstract

Regulation of RNA splicing is pivotal in neural development, with established gene isoform expression patterns (*1, 2*). However, the specific roles of isoform diversity across cell types in both healthy and diseased brains warrant further investigation. Here, we employed a combination of metabolic RNA labeling using 4-thiouridine (4sU) and long-read sequencing based single-cell full-length transcriptome sequencing to capture newly synthesized transcripts within the developing mouse cortices. This approach allowed us to identify predetermined cell states supported by new RNAs and the driving isoforms of transcription factors that regulate the development of deep-and upper-layer neurons. Through detailed single-cell isoform expression analysis, we discovered novel cell type-specific isoforms and uncovered isoform switch events that modulate neuron differentiation. Additionally, we investigated isoform regulation associated with Autism Spectrum Disorder (ASD) during embryonic development of BTBR *T*+ *Itpr3tf* (BTBR) mice. Notably, our findings indicate a premature emergence of cortical projection neurons (CPNs) with an immature identity in ASD-affected cortices. These CPNs exhibit the highest degree of differential transcript usage (DTU), significantly overlapping with their cell type markers and being enriched in RNA splicing processes. Exon inclusion was significantly enriched in ASD and the related RNA binding proteins (RBPs) were extracted, nearly 60% of which have been reported as ASD risk genes. Lastly, we revealed a reduction in isoform diversity in ASD, potentially linked to H3K27ac dysregulation in the associated genes. Collectively, our study marks a substantial advancement in understanding the molecular basis of cortical development and function, paving the way for future studies on neurodevelopmental disorders.

## INTRODUCTION

The cerebral cortex, a highly intricate structure, plays a central role in mammalian brain function and undergoes sophisticated developmental processes that encompass orchestrated gene expression and cell lineage diversification. Recent advancements in single-cell RNA sequencing (scRNA-seq) have significantly enhanced our comprehension of cortical development, offering an unprecedented level of resolution to delineate cellular heterogeneity and transcriptional alterations in both murine and human models. Utilizing scRNA-seq in mice, researchers have generated detailed transcriptional roadmaps of cortical development, uncovering dynamic gene expression patterns across various cell types during these processes (*3–8*). In the human context, the integration of transcriptomics, epigenomics, and chromatin state data have offered a comprehensive view of detailed single-cell resolution map of human cortical development, shedding light on the gene regulatory mechanisms, cellular diversity, and epigenetic dynamics that shape the human brain during its early development (*9–18*).

While these studies have offered significant insights into the transcriptional landscapes of cortical development, the role of alternative splicing, a key mechanism for expanding transcriptome diversity, remains underexplored due to inherent limitations of short-read sequencing based scRNA-seq. Alternative splicing plays pivotal roles in brain development, critically influencing neuronal differentiation, synaptogenesis, and the stratification of cortical layers (*19, 20*). For example, synaptic genes such as *Nrnx1* have been found to harbor thousands of unique isoforms (*21*). Additionally, the isoform repertoire of protocadherin, beyond the three γC-type isoforms, is essential for neuronal survival, underscoring the isoform-specific functions in neurodevelopment (*22*). In contrast to conventional short-read sequencing, the third generation sequencing (TGS) directly captures the complete RNA sequence leveraging its long read length advantage, providing a detailed blueprint of alternative splicing and transcript architecture. Long-read sequencing studies in mice have unveiled neuron-and glia-specific splicing patterns during both development and adulthood, emphasizing the functional significance of splicing regulation in the diversification of cortical cell lineages (*23*). Moreover, such studies in the developing human brain have uncovered the intricate splicing patterns that are central to brain development and disease risk (*24*). However, the application of isoform-level data to reconstruct developmental trajectories in the cortex or to analyze transcript-specific changes along differentiation pathways remains sparse.

Most RNA-seq approaches provide only a static snapshot of the total RNA pool within the cell, offering little information on the temporal transcription activities because of a mixture of newly transcribed (‘new’) and pre-existing (‘old’) RNAs. Several methods such as scSLAM-seq and scNT-seq have been devised to distinguish new RNAs and old RNAs within single cells (*25*). RNA velocity, a powerful tool for probing dynamic cellular processes, is often constrained by assumptions regarding the distribution of spliced and unspliced RNA populations. The Incorporation of 4-thiouridine (4sU) labeling for newly synthesized RNA offers a refined approach to enhance RNA velocity analysis (*26*). By directly tagging nascent RNA with 4sU during transcription, we observed a clear demarcation between new and old RNAs, thereby mitigating biases introduced by transcriptional or degradation dynamics. When integrated with TGS technologies, such as PacBio or Nanopore, 4sU labeling enhances the resolution of nascent transcript detection, capturing full-length RNA molecules and complex splicing events. This synergistic approach provides an unparalleled framework for deciphering the transcriptional dynamics that underpin cellular differentiation and lineage decisions, offering profound insights into cell fate transitions and developmental processes.

Autism spectrum disorders (ASD), classified as a neurodevelopmental disease (*27*), are deeply rooted in atypical brain development, particularly affecting the structure and function of the nervous system. The hallmark symptoms of autism, emerging during early childhood (around ages 2–3), are primarily associated with social interaction, communication, and behavior. A burgeoning body of studies has implied the development of ASD from embryonic neurogenesis. ASD patients-derived organoids exhibit perturbed neurodevelopmental process, characterized by an accelerated cell cycle and overproduction of GABAergic neurons (*28*), and have revealed dorsal intermediate progenitors, ventral progenitors and upper-layer excitatory neurons as the most vulnerable cell types (*29, 30*). Animal models have further shed light on these mechanisms. CHD8 conditional knockout mouse models, for instance, exhibit increased cortical thickness and reduced synaptic plasticity (*31*), neuroanatomical changes that are associated with behavioral deficits reminiscent of ASD, such as impaired social interactions. The investigations of human organoids and animal models underscores the intricate interplay between early neurogenesis and ASD pathology. However, the absence of diagnostic methods for patients during the embryonic stage precludes the study of ASD during embryonic development. BTBR T+ Itpr3tf (BTBR) mouse, an inbred strain exhibiting multiple ASD-like behavioral phenotypes, is commonly utilized to investigate the underlying mechanisms of ASD and to explore therapeutic potential (*32–35*). Emerging evidence implicates alternative splicing as a critical mechanism linking gene variation and neuropsychiatric disease (*36–41*). Yet how isoform expression shifts during the embryonic stages of ASD candidates, particularly at cell-type resolution has never been investigated.

In this study, we employed TGS-based single-cell full-length isoform sequencing in developing mouse cerebral cortices, seeking to identify the driving TF isoforms for neuron specification by combining with new RNA labeling. More importantly, we compared isoform diversity and isoform switch regulation in normal and diseased cortical development, identifying the ASD-related splicing events and RNA binding proteins (RBPs). Our work provides a comprehensive view of the complexity of full-length transcripts in neural development at single-cell resolution, with critical implications for understanding regulatory mechanisms of neuron fate specification and identifying risk factors of ASD.

## RESULTS

### Cell type identification and transcriptional profiling in developing cerebral cortex

We generated comprehensive single-cell expression profiles at both the gene and isoform levels throughout the developmental stages E13.5, E15.5, and E17.5 of mouse cortex, employing the next generation sequencing (NGS)-based short-read sequencing and TGS-based long-read sequencing simultaneously (Fig. 1A). After quality control, our dataset comprised 55,773 cells with the gene expression profiles (averaging 2069 genes per cell) and 12,418 cells with the isoform expression profiles (averaging 837 isoforms per cell), the latter being attributed to the constrained sequencing depth of TGS (fig. S1A and S1B). To check whether the low depth of TGS impacted the fidelity of gene expression profiling, we conducted a comparative analysis of the two datasets at both the single-cell and bulk levels. The correlation analysis revealed a positive correlation between the gene counts and unique molecular identifier (UMI) counts from the two platforms, with coefficients of 0.82 and 0.79, respectively (fig. S1C). Principle component analysis (PCA) demonstrated that despite technical batch variations (PC1 in fig. S1D), they both datasets effectively captured the molecular distinctions between different developmental stages (PC2 in fig. S1D).

**Fig. 1.**
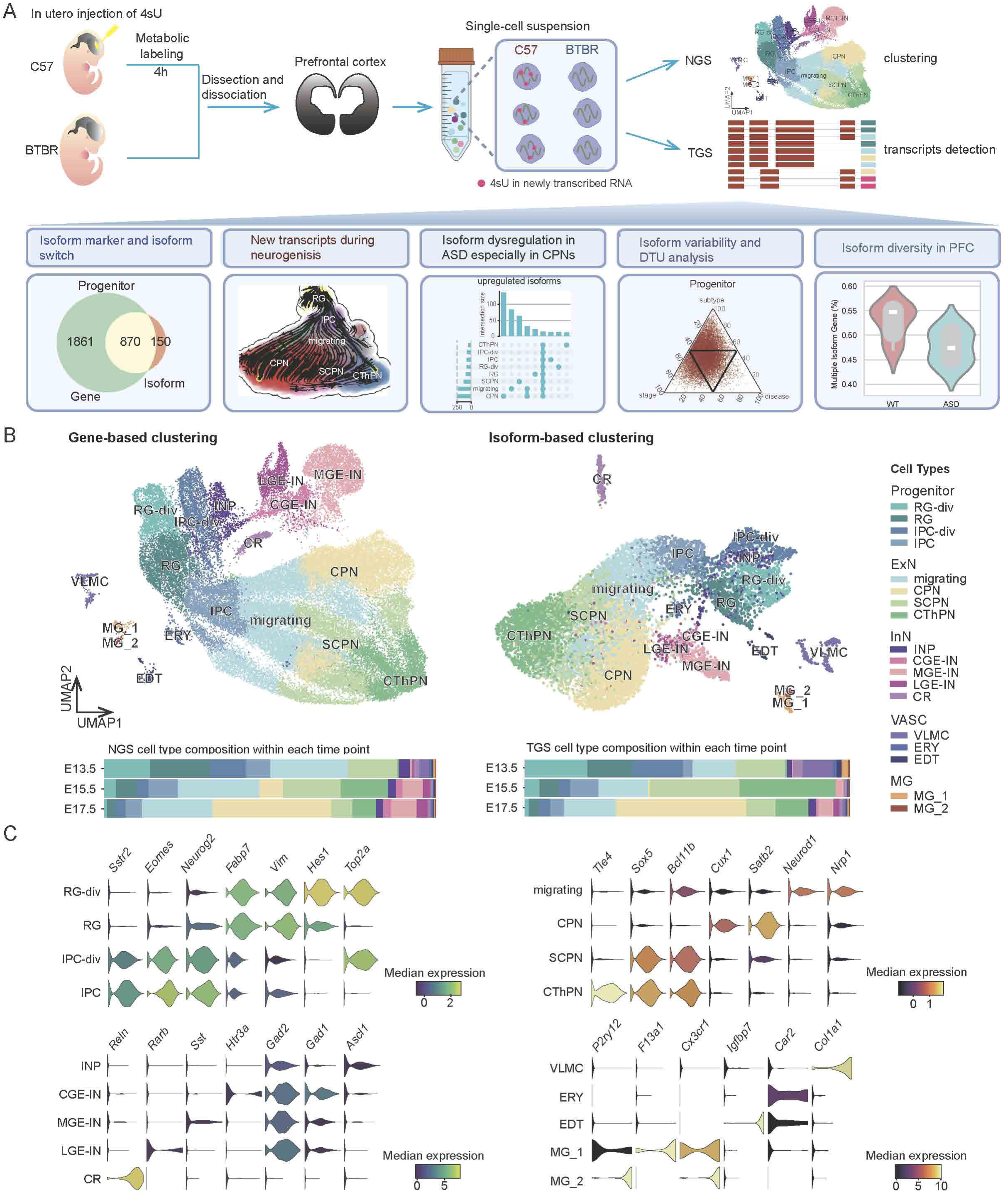
Overview of the study and cell type analyses in the developing mouse cerebral cortices. (**A**) Experimental design for gene and isoform profiling of the developing mouse prefrontal cortex at single-cell resolution combined with new RNA labeling. (**B**) Uniform manifold approximation and projection (UMAP) embedding of cells with the NGS-detected gene expressions (left) and with the TGS-detected isoform expressions (right) from E13.5, E 15.5 and E17.5 mice. RG, radial glia; RG-div, dividing radial glia; IPC, intermediate progenitor cell; IPC-div, dividing intermediate progenitor cell; migrating, migrating neuron; CPN, callosal projection neuron; SCPN, subcerebral projection neuron; CThPN, corticothalamic projection neuron; MGE-IN, medial ganglionic eminence-derived interneuron; CGE-IN, caudal ganglionic eminence-derived interneuron; LGE-IN, lateral ganglionic eminence-derived interneuron; CR, Cajal–Retzius cell; VLMC, vascular leptomeningeal cell; EDT, endothelial cell; ERY, erythrocyte cell; MG, microglial cell. (**C**) Violin plot showing the expression levels of representative marker genes per cell subtype in the NGS dataset.

Upon further assessment of the efficiency of restoring different cell types in both datasets, it was observed that, despite capturing less than a third of the cell sample size of NGS, TGS identified the same five major cell types, encompassing neural progenitors (Progenitor), excitatory neurons (ExN), inhibitory neurons (InN), microglias (MG) and vascular cells (VASC) (Fig. 1B). These cell types showed distinct expression patterns at both the gene and isoform levels (fig. S2A, S2B). The major cell types were further clustered into 18 subtypes, each characterized by the expression of representative marker genes (Fig. 1C). Hierarchical clustering of all cell subtypes at both gene and isoform levels revealed similar relationships (fig. S1E). Notably, neuronal subtypes (except for the CR cells) were more closely related to each other and showed the greatest divergence from non-neuron subtypes. Meanwhile, erythrocyte cells exhibited larger distance from other cell types in gene expression measured the TGS, indicating a higher resolution in distinguishing between cell types with TGS-based profiling (fig. S1E).

We also evaluated the cellular composition at each developmental stage (Fig. 1B). As expected, in both technical datasets, the proportion of progenitors was highest in the earliest stage and progressively declined along development (∼50% at E13.5, ∼22% at E15.5, ∼11% at E17.5, Fig. 1B). The upper-layer neurons (CPN) were sparse at E15.5 and largely increased in E17.5, suggesting the temporal development of distinct neuron types. The temporal dynamics of cell type distribution were consistent across both NGS-based gene profiles and TGS-based isoform profiles (fig. S1F). In summary, our approach accurately captured the cellular landscape along with the corresponding gene and isoform expressions in the developing mouse cerebral cortices.

### Isoform switch and diversity in neuronal development

We extracted cell type markers at gene and isoform levels respectively (table S1), and compared the consistency. Due to the larger dataset of NGS, each cell type was detected with more differentially expressed genes (DEGs) than differentially expressed isoforms (DEIs) (Fig. 2A). The majority of DEIs belong to the DEGs for each cell type, indicating the consistency of both datasets in capturing the molecular characters. Meanwhile, there were some genes only appearing as DEIs, indicating switches of isoform expression being masked at gene level. By further comparing the isoform expressions between Progenitor and ExN, we identified pronounced isoform switching for 13 genes, including those being reported to involve in neuron development and brain functions, such as *Clta* (*42, 43*)*, Gpm6b* (*44*)*, etc* (Fig. 2B). The isoform switching events were classified into two types: classical switches, where the dominant isoform alternates, and non-classical switches, where the dominant isoform remains consistent (Fig. 2C). For example, *Clta,* which has been reported as a classical switch example (*45*), was also captured in our data (Fig. 2D). *Ergic3*, which belongs to the COPII-associated proteins (*46*), showed dominant expression of *Ergic3-201* in Progenitor but *Ergic3-202* isoform in ExN (Fig. 2D). *Tmod3* is an example of non-classical switch where *Tmod3-201* down-regulated while *Tmod3-205* up-regulated from Progenitor to ExN. We further verified these isoform switch events with reverse transcription and PCR (fig. S2C). The identification of these isoform dynamics provides novel insights into the molecular underpinnings of neurodevelopment and suggests a potential layer of regulation that may be critical for the precise regulation of neurogenesis.

**Fig. 2.**
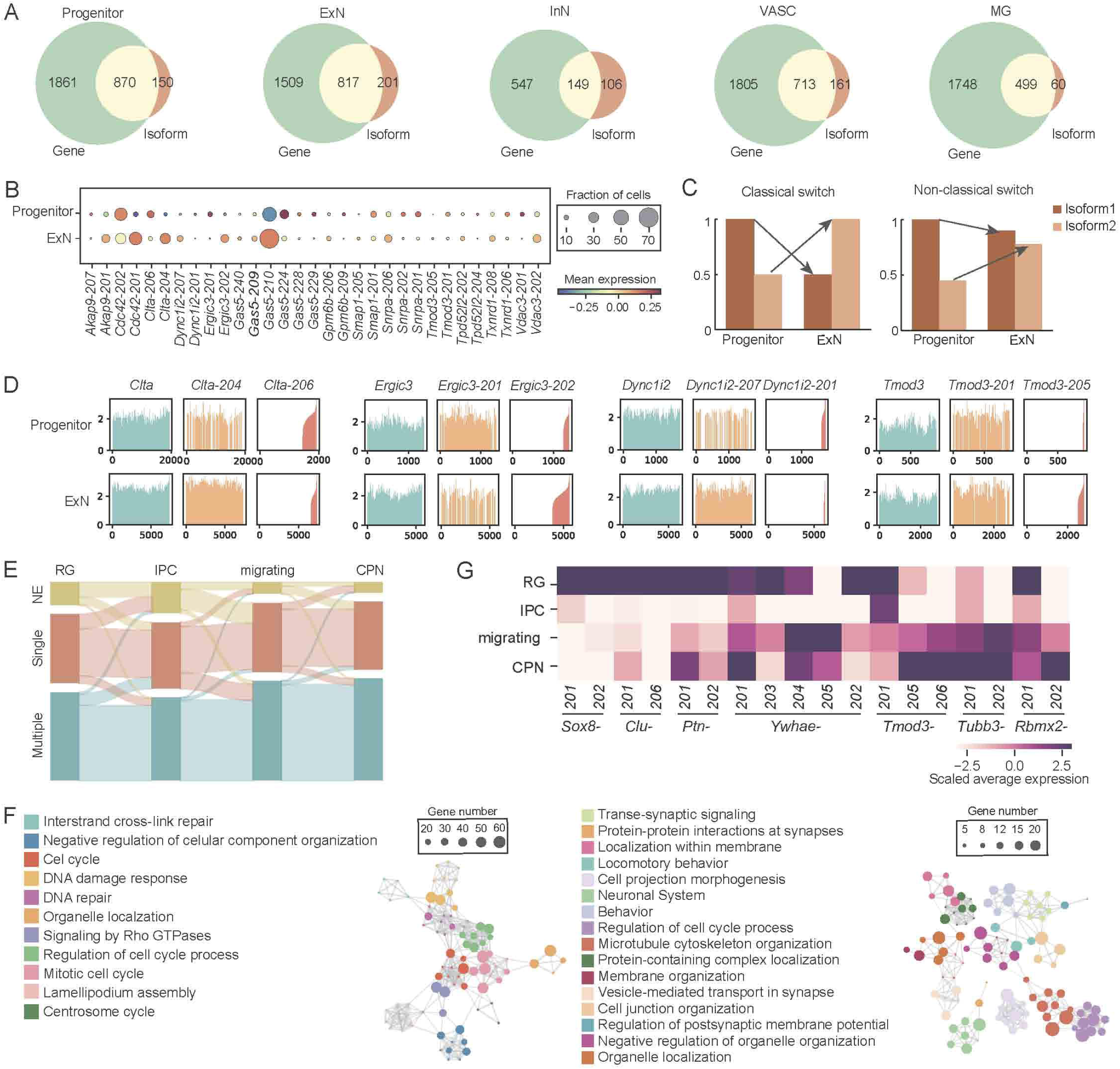
Isoform switch and diversity in neuronal development. **(A)**Venn diagrams represent the overlap of cell type DEGs (green) and DEI related genes (brown). **(B)** Dotplot showing the scaled isoform expression values of top 13 genes showing isoform switch in Progenitor and ExN. **(C)** Representative cartoons exemplify the isoform expression patterns of the two types of isoform switch. **(D)** Bar plot showing the single-cell expression levels of indicated genes and isoforms in Progenitor and in ExN. **(E)** Sankey diagram showing the dynamic changes of isoform diversity for each gene along cortical neuron development from RG to CPN. **(F)** Enrichment network representing the top enriched biological terms of genes with multiple expressed isoforms in RG but single expressed isoform or not expressed in IPC (left) and genes switched to multiple isoforms expression from IPC to migrating neuron (right). Each dot represents a gene ontology (GO) term and the size represents the number of hit genes in the term. GO terms belonging to the same categories are labeled with the same color and each pair of terms are connected by a line if they share common genes.. **(G)** Heatmap showing the examples of isoform diversity changes across different cell types in CPN neuron development.

Furthermore, we assessed isoform diversity across various cell types related to excitatory neuron development. A total of 13300 genes were detected in all cells, which were divided into three categories in each cell type: non-detected expression (NE), detected with single isoform expression and detected with multiple isoform expression (table S2). During excitatory neurogenesis, the isoform diversity transiently reduced from RGs to IPCs, with more NE and single isoform genes in the latter (Fig. 2E). GO analysis revealed these genes were almost related to cell cycle (Fig. 2F). For instance, *Sox8,* which is essential in maintaining and promoting the proliferation of neural stem cells (*47*), exhibited high expression levels of two isoforms in RG but low expression levels of only *Sox8-201* in IPC (Fig. 2G). Then more genes activated expression or generated multiple isoforms during IPC to migrating neuron development, such as *Tubb3* and *Rbmx2* (Fig. 2E, 2G). These genes were enriched in diverse biological processes, including cell projection morphogenesis, neural system, behavior, localization within membrane, etc. (Fig. 2F). For example, *Ywhae*, also known as *14-3-3ε* (Fig. 2G), plays a crucial role in various aspects of neural development, including neuronal migration, cytoskeletal organization, neurogenesis, and synaptic function (*48*). During further lineage specification (to CPN, SCPN and CThPN), the isoform diversity all showed few changes (Fig. 2E, fig. S2D). About 40% of the RG-initiated multi-isoform genes maintained transcript diversity until the projection neurons (Fig. 2E), indicating the sustained requirement for these isoforms function throughout neurodevelopment. The increase in isoform diversity underscores the complexity of the mature neuronal transcriptome and the potential for isoform-specific regulation of neuronal function.

### Nascent isoform expression revealed driving TF isoforms for neuron specification

To capture the crucial RNA dynamics regulating *in vivo* neural development, we labeled the embryonic cerebral cortices in utero with 4sU for four hours and then collected the cells for full-length isoform sequencing (Fig. 1A). The NGS-based scRNA-seq only detected a small fraction of each RNA molecule, which may lead to underestimation of the amount and proportion of newborn RNAs in a process (Fig. 3A). Our full-length RNA sequencing shall capture the new RNAs more efficiently. We firstly checked the labeling efficiency of our experimental approach. All samples showed specific T-to-C substitution as expected, with no difference between the NGS and TGS platforms for each sample (fig. S3A). The T-to-C substitution rate was around 2% for all samples except the E17.5-2 replicate showed aberrant high labeling efficiency (∼8% T-to-C substitution rate). As expected, the detected new to total RNA ratio (NTR) in the TGS dataset was much higher than that in the NGS dataset, both measured by cells and genes (fig. S3B). The higher labeling rate of E17.5-2 leads to higher NTR compared with E17.5-1. To determine whether this difference affected the expression profiles, we calculated the expression correlation between the two E17.5 replicates in both old and new RNAs. Both correlation values were high, with 0.95 for old RNA and 0.96 for new RNA (fig. S3C). Thus our single-cell new RNA sequencing procedure was reliable and the results were repeatable.

**Fig. 3.**
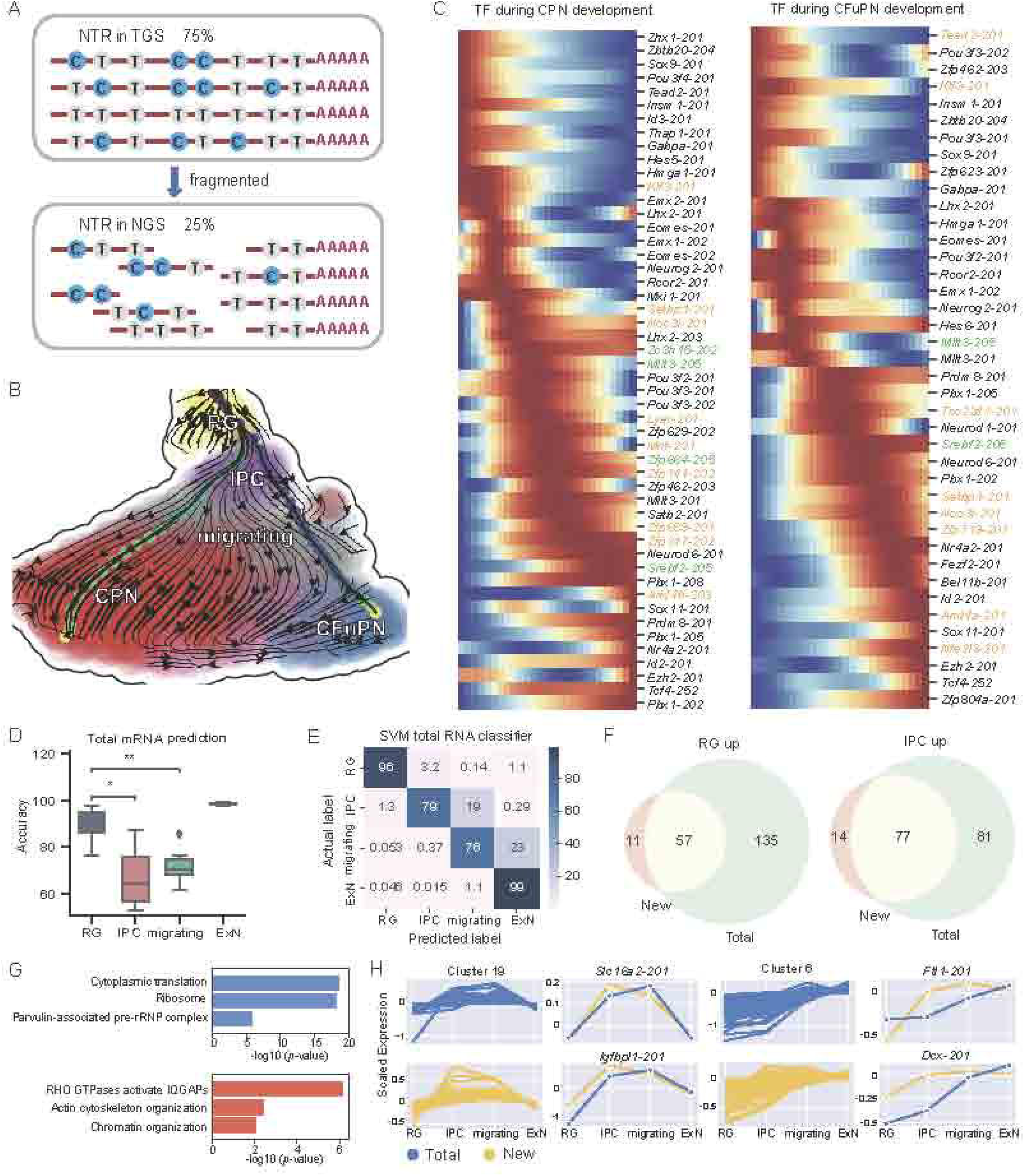
Total and new transcript analysis in neurodevelopment. (**A)** A schema depicting more precise NTR detection by our TGS-based full-length isoform sequencing.. (**B**) RNA velocity constructed by *Dynamo*, based on 4sU metabolic isoform expression. The trajectory illustrates two sublineages: upper-layer (CPN) neurons and deep-layer (CFuPN, including CThPN and SCPN) neurons, both originating from RGs. (**C**) Regulatory TF isoforms along the CPN and CFuPN lineages. The pseudotime was divided into 27 bins, and isoforms were ordered by the pseudotime of their maximum expression along the trajectories. Isoforms which have not been reported are highlighted in yellow, non-coding isoforms whose genes were reported as functional in the respective lineage are in green. (**D**) Boxplot showing accuracy of the six machine learning classification methods (SVM, RF, LR, MLP, GB, DT) in predicting cell type labels using new RNA expression profiles. Mann-Whitney Wilcoxon test was conducted for the pairs RG-IPC, RG-migrating \**p* < 0.05, \*\**p* < 0.01. (**E**) Confusion matrix showing the accuracy of SVM. (**F**) Venn Diagram showing the number of DEIs between RG and IPC cells calculated by new and total RNAs. (**G**) Represent GO terms of RG upregulated isoforms (top) and IPC upregulated isoforms (bottom) specific in new RNA analysis. (**H)** Representative clusters of genes showed dynamic expression changes from RG to ExN development. Example isoforms showing unmatched changes in the two type of RNA profiles are shown.

Then, we analyzed developmental trajectories of excitatory neurons using the new RNA information by dynamo (*25*). There were two branches sharing the same origin in RG but immediately separated in the IPC stage, then specified into upper-layer neurons (CPN) and deep-layer neurons (CFuPN) (Fig. 3B). The fate diverging nodes inferred from new RNAs supposed to be earlier than total RNA based analysis, where the neuron fate differentiation was observed in the migrating neurons (*3*). Then we exacted the key TF isoforms regulating RG to CPN and RG to CFuPN, respectively (Fig. 3C) (*15*). Most of the TFs have been reported to play crucial roles in neurodevelopment, such as *Sox9* (*49*), *Neurog2* (*50*), *Satb2* (*51*), *Fezf2* (*52*). We accurately specified the exact isoforms of these genes function in the process. Moreover, we identified the functional roles of different isoforms of the same gene during neuron lineage specification. For example, *Lhx2* is known to play critical roles in the migration and projection of postmitotic neurons that form the upper layer of the cerebral cortex during neurogenesis (*53*). However, the roles of different isoforms has not been well-documented. According to our study, *Lhx2-201* was mainly distributed in early CPN lineage and CFuPN lineage while *Lhx-203* was highly expressed in CPN lineage from migrating to CPNs. These findings highlight the importance of isoform resolution in understanding the molecular logic of neuronal lineage progression.

### Cell fate is predetermined according to new RNA analysis

Our investigation into the precocious determination of cell fate, as inferred from the analysis of new RNA profiles, revealed intriguing discrepancies between new RNA and total RNA pool. To elucidate this divergence, we conducted a comparative analysis of the RNA profiles from both perspectives (Fig. 3D). Firstly, we trained a suite of six machine learning classifiers-encompassing Random Forest (RF), Multilayer perceptron (MLP), Support Vector Machine (SVM), Decision Tree (DT), GradientBoost (GB) and Logistic Regression (LR), to discern cell type labels using the comprehensive total mRNA profiles. These classifiers were then utilized to predict cell type labels based on the new mRNA profiles. It revealed a high concordance for ExNs but a significant discrepancy for IPCs and migrating neurons, with less than 80% accuracy in classification (Fig. 3D). Intriguingly, a subset of IPCs (ranging from 9.4% to 37%) were misidentified as migrating neurons, and a substantial proportion of migrating neurons (17% to 35%) were incorrectly classified as ExNs based on new mRNA profiles (Fig. 3E, fig. S4A). This finding suggests that the new mRNA profiles of IPCs and migrating neurons closely resemble the total mRNA compartment of the subsequent cell stage.

To substantiate the asynchronous nature of the total and new RNA profiles, we performed dimensionality reduction analysis on merged datasets. The analysis revealed a homogeneous mixture of total and new RNA profiles in RG, with a clear divergence at the IPC stage (fig. S4B, C). To account for potential biases due to sequencing coverage, we down-sampled the total RNA to 20% of its original depth, mirroring the NTR. Despite this adjustment, the distinction between total and new RNA profiles persisted at the IPC stage, indicating that the observed differences were not an artifact of sequencing depth (fig. S4D). Given that the new RNA profile of IPCs already encapsulated the RNA signatures of migrating neurons, we hypothesize that the divergence of the two RNA profiles may be attributed to the onset of neuronal differentiation.

To uncover the molecular underpinnings driving the divergence between new and total mRNA profiles at the onset of neural differentiation, we extracted DEIs between RGs and IPCs within both RNA profiles. We identified 11 and 14 isoforms specifically upregulated in the new RNAs of RG and IPC, respectively (Fig. 3E). The 11 new RNA-specific DEIs in RG were predominantly ribosomal protein genes (Rpl and Rps), consistent with the high cellular proliferation, metabolic demands, and protein synthesis requirements of stem cells (Fig. 3G, fig. S4E, table S3). In contrast, GO analysis of the 14 new RNA-specific DEIs in IPC revealed RHO GTPase activation and actin cytoskeleton organization, such as *Actb-201* and *Tubb5-201* (Fig. 3G, fig. S4E, table S3). This suggests that IPCs possess an enhanced capacity for migration and motility.

To systematically identify isoforms with distinct expression patterns during neurogenesis, we applied k-means clustering to categorize the expression trends of both new and total mRNA isoforms. Notably, cluster 19 represented isoforms that were transiently activated in the intermediate cell types, while cluster 6 showcased isoforms that were consistently upregulated from RG to ExN (Fig. 3H). Although the expression patterns of the majority of isoforms were congruent between the two RNA profiles (such as *Neurod1-201* and *Mef2c-222* in fig. S4F), certain genes related to neuronal development displayed obvious differences. For example, the expression of new RNA for *Slc16a2-201* (*54*) and *Igfbpl1-201* (*55*) peaked in IPCs, whereas the total RNA for the two isoforms exhibited the highest expression in migrating neurons. Similarly, *Ftl1-201* and *Dcx-201* in cluster 6 showed markedly elevated new RNA expression levels in IPCs. The observation that some nascent transcription is most active in earlier cell types suggests that these pathways may be initiated earlier in neurodevelopment than previously observed. This early transcriptional activation may precipitate critical neurodevelopmental events, laying the groundwork for subsequent differentiation and migration processes in neurons.

### CPN neurons are largely affected in ASD pathogenesis

To ascertain the presence of aberrant gene and isoform expressions at the embryonic stage in ASD models, we conducted a comprehensive profiling of single-cell full-length transcripts in the developing cortices of BTBR mice. Our analysis encompassed 19,059 cells profiled using the NGS platform and 4,602 cells profiled using the TGS platform, with a median of 2,192 genes and 964 transcripts per cell. The cell types identified in ASD samples were consistent with those in wildtype (WT) mice. (Fig. 4A, 4B, fig. S5A and S5B). The majority of the DEGs between WT and ASD were common across cell types, indicating a pervasive dysregulation, with the exception of CPNs that exhibited a higher number of cell type-specific DEGs ( fig. S5C, table S4). This observation aligns with existing literature, highlighting the complex gene regulation patterns in CPNs during development and their increased susceptibility to neurodevelopmental disorders such as ASD (*41, 56, 57*).

**Fig. 4.**
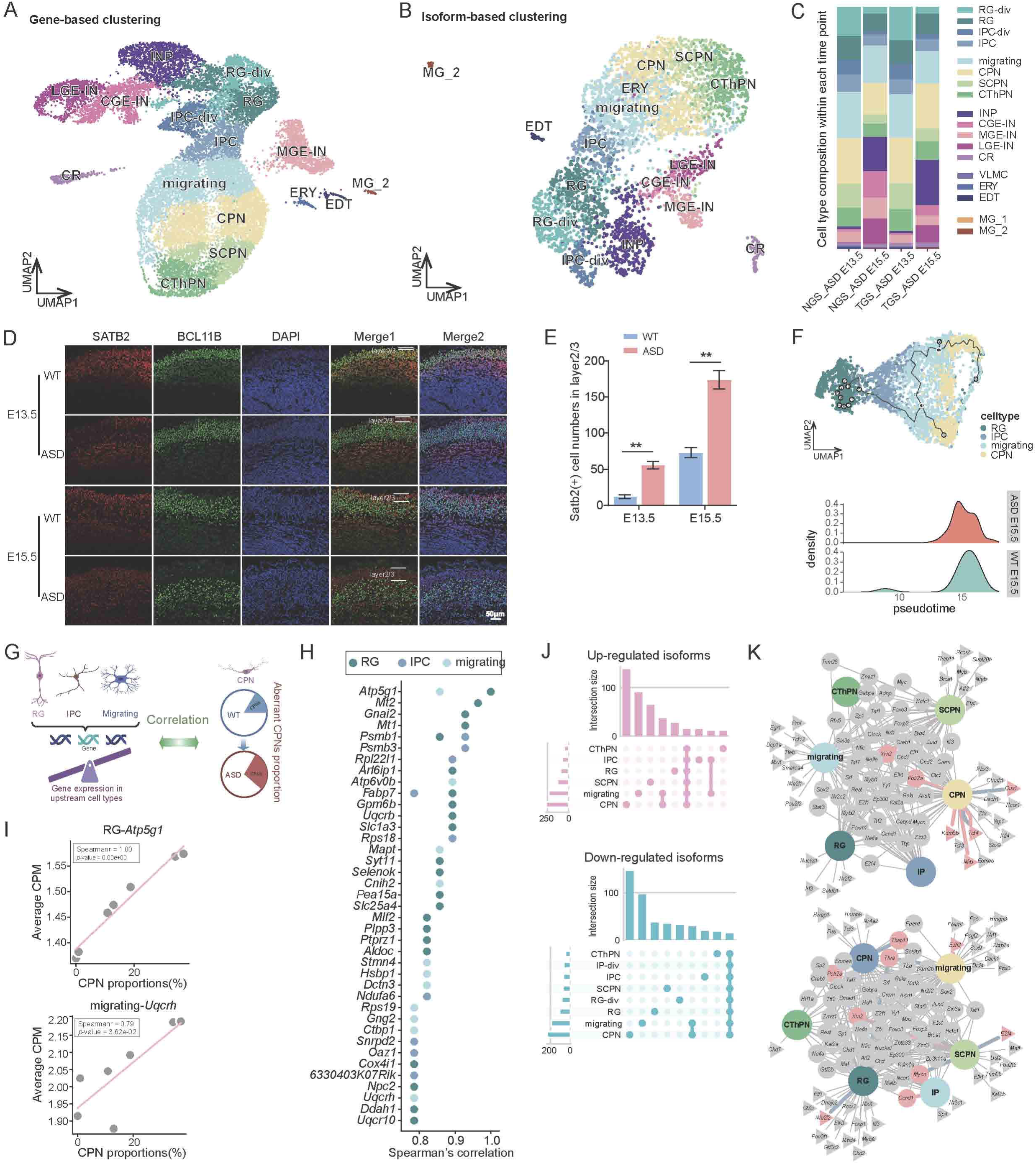
Pre-development of CPNs in ASD. (**A-B**) UMAP showing the cell types from developing cortices of BTBR ASD model based on gene expression (**A**) and isoform expression (**B**). (**C**) Cell type proportions in each developing stage of ASD model. (**D**) Immunostaining of the CPN marker SATB2 and the deep-layer neuron marker BCL11B in WT and BTBR cortices; n = 3 per group. Scale bars: 50 μm. (**E**) Cell counts of SATB2 positive neurons in layer 2/3 (per 45 mm^2^) according to the immunofluorescence results in panel 4D. \*\**p* < 0.01. (**F**) Developmental trajectory with cell types colored (top) and pseudotime distribution of ASD CPNs and WT CPNs respectively (bottom) (**G**) Outline of the analysis linking gene expression of RG, IPC and migrating neuron to the proportion of CPN in WT and ASD mice. (**H**) Spearman’s correlation coefficients of expression levels of the 39 “CPN predictor genes” within each progenitor cell type and the proportions of CPN (Spearman correlation coefficients > 0 and p < 0.05) (**I**) Scatter plot of CPN proportions and averaged expression of *Atp5g1* in RG cells and *Uqcrh* in migrating neurons respectively. Each dot represents one sample. (**J**) Upset plot showing cell type specific and common DEI numbers between WT and ASD. (**K**) Cell type-TF networks predicted by ASD-upregulated (top) and downregulated (bottom) DEGs. Genes in circle nodes represent they are regulated in at least two cell types. Triangular nodes represent cell type-specific regulators. Gray nodes represent regulators themselves not differentially expressed and red nodes represent regulators themselves were detected as DEIs. Red lines represent the regulators upregulated in ASD and blue lines represent those downregulated in ASD.

We further compared the DEGs to previous datasets related to ASD study, including those from ASD patients (*58*) and adult ASD mice (*59, 60*) (fig. S5D). The DEGs in each cell type were significantly enriched in ASD-associated genes identified by M. Wang et al., who investigated the young ASD mice (fig. S5E). Moreover, these DEGs were significantly enriched in matched cell types from ASD organoids (*30, 61*), particularly in excitatory neurons (fig. S5F). In contrast, none of these DEG sets were enriched in the ASD genes identified from the adult brains (S Mizuno et al., MJ Gandal et al., and SFARI in fig. S5E). This result underscores a relatively lower concordance of the ASD-related gene features between embryonic and adult stages, emphasizing the importance of studying the brain during developmental stages.

Upon examining the cellular composition of ASD cortices during neural development, we unexpectedly observed large abundances of CPNs in E13.5 ASD cortices, contrasting with their emergence from E15.5 in WT cortices (Fig. 4C). This premature emergence of CPNs in ASD was further validated through immunofluorescence, which revealed a significant increase in SATB2+ cells in layer 2/3 (Fig. 4D, E). Pseudotime analysis also indicated that E15.5 ASD CPNs exhibit a less mature state compared to their WT counterparts, with the majority of ASD CPNs positioned at earlier pseudotime stages (Fig. 4F, fig. S6A). This finding suggests a disruption in neurodevelopmental timing, potentially contributing to the altered developmental characteristics observed in ASD.

Given that CPNs are terminal cells of neurogenesis, the abnormal proportion of these cells may be linked to dysregulated progenitor populations. We conducted a correlation analysis to identify causal genes in progenitor cells associated with variations in CPN proportions (Fig. 4G) (*30*). We identified 39 genes whose expression levels in upstream cell types were positively correlated with increased CPN proportion (Fig. 4H). The majority of these genes are known regulators involved in neurodevelopment, implicated in processes such as energy metabolism, synaptic function, or protein synthesis (for example, *Fabp7*, *Gpm6b* and *Aldoc*). Meanwhile, there were less characterized TFs such as *Oaz1* and *Npc2*. *Atp5g1*, a subunit of ATP synthase, is vital for mitochondrial ATP production, and its role in RG is vital for supporting the high-energy demands of neurogenesis, neuronal migration, and differentiation (*62*). *Uqcrh* (Ubiquinol-Cytochrome c Reductase Hinge Protein), as part of complex III, has been shown to lead to various neurodevelopmental disorders (*63*), including ASD, intellectual disabilities, and neurodegenerative conditions. The expression level of *Atp5g1* in RGs, and of *Uqcrh* in migrating neurons were positively correlated with CPN proportion, respectively (Fig. 4I).

### Aberrant isoform expression in ASD during embryonic development

Expanding our analyses to the transcript expression level, we observed a widespread pattern of DEIs in the developing cortices of the ASD model (Fig. 4J, table S4). Consistent with the patterns observed in DEGs, CPNs displayed the most pronounced differences at the isoform level. Among the top dysregulated isoforms in CPNs, we identified genes pivotal for synaptic function, including *Idh3b-202* and *chd3-201*, as well as the TFs *Nnat-205* and *Itb2b-201* (*64*). To substantiate the DEI results, we conducted RT-qPCR on 12 transcripts (fig. S6B). Collectively, our data indicate a disruption in developmental and synaptic signaling within components of upper-layer cortical circuitry in ASD. Both upregulated DEGs and DEIs in ASD showed enrichment in pathways associated with mTORC1 signaling and cell cycle regulation, including E2F targets and G2-M checkpoint (fig. S6C), aligning with previous ASD research (*58*). Wang et al. (*65*) have highlighted mitochondrial dysfunction (e.g., oxidative phosphorylation) as a recurring theme in ASD mouse models and human datasets, contributing to abnormal neuronal growth and metabolism. In our dataset, oxidative phosphorylation enrichment was observed exclusively in DEI analysis, suggesting that splicing regulation may modulate these pathways during embryonic stages.

To unravel the regulatory mechanisms, we inferred upstream TFs associated with ASD DEIs. We identified 89 TFs for the upregulated ASD DEIs and 97 TFs for the downregulated ASD DEIs, with 15 of these TFs also being ASD DEIs themselves (Fig.4K, table S4). TFs such as *Myc*, *E2f1*, *Foxm1* were implicated in increasing isoform expressions across multiple cell types in ASD, reflecting their broad influence in cell growth and proliferation pathways, which are known to be hyperactive in ASD models (*56, 66*). Certain TFs, including *Nelfa* and *Kat2a*, were suggested to specifically regulate DEIs in RGs and CPNs respectively, indicating that transcriptional changes in the ASD model are finely tuned within specific neuronal populations, potentially contributing to cell type-specific vulnerabilities observed in ASD. Notably, an isoform switch event in the *Tcf4* gene in ASD CPNs was characterized by the upregulation of the non-coding isoform *Tcf4-211* in ASD, while the coding isoform *Tcf4-252* was downregulated (fig. S6B, S6C). Transcription Factor 4 (TCF4), a helix-loop-helix (bHLH) transcription factor, which was required for correct brain development during embryogenesis and has been associated with autism, schizophrenia, and other neuropsychiatric disorders (*67–71*). This shift in our study may suggest a potential mechanism of isoform regulation, where increased levels of non-protein-coding mRNAs could competitively inhibit the expression of protein-coding mRNAs. While these changes could be obscured by overall mRNA levels, they may be reflected in the protein abundance of *Tcf4*. The identification of additional isoform switch events helps explain the enrichment of the same signaling pathways in both upregulated and downregulated DEIs in ASD (fig. S6D, S6E). These findings underscore the critical role of isoform-level regulation in the complex pathophysiology of ASD.

### Pseudogenes are commonly dysregulated across cell types in ASD

Pseudogenes, traditionally regarded as non-functional DNA sequences arising from the nonsense mutations or disruptions of protein-coding genes, have emerged as potential players in modulating complex phenotypes, including those associated with neurodevelopmental disorders such as ASD (*72, 73*). However, the functional annotation of pseudogenes, particularly within the context of neurodevelopment disease, has been constrained by the limitations of NGS-based scRNA-seq. Leveraging the ability to obtain full-length transcripts, we accurately differentiated pseudogenes from their parent genes, revealing a significant proportion (12.9%) of common DEIs that were pseudogenes, in contrast to the 1.4% observed in cell-type-specific DEIs (fig. S6F). This observation suggests that pseudogenes may broadly influence transcriptional regulation or serve as a reflection of the cellular response to the perturbed environment in ASD.

Among the pseudogenes identified, the isoform *2610005L07Rik-202 was a* common DEI (fig. S6G, table S4), with its parent gene being *Cdh11.* Previous research has established a link between the absence of Cdh11 and the manifestation of multiple autistic-like behaviors in mice (*74–76*). Furthermore, iPSC-derived cortical organoids from individuals with ASD have shown downregulation of *Cdh11* (*77*), which was also presented in our data (fig. S6H). Pseudogenes are hypothesized to function as competitive endogenous RNAs (*78*), sponging microRNAs (*79*) and thereby modulating the expression of their target genes. To elucidate the regulatory relationship between pseudogenes and their parent genes, we overexpressed *2610005L07Rik-202* in cultured neurons derived from mouse embryonic cerebral cortices. Interestingly, contrary to our expectations, *Cdh11* showed a little bit of upregulation after overexpression of *2610005L07Rik-202* (fig. S6I). This finding indicates that the regulatory interplay between pseudogenes and their parent genes is complex and may involve distinct mechanisms *in vivo* and *in vitro*.

### Perturbed isoform switching events and AS-related RBP regulators in ASD

To explore the functional implications of splicing programs across cell subtypes, developmental stages and neurodevelopmental diseases, we comprehensively analyzed isoform variability across these critical dimensions (fig. S7A) (*23*). For Progenitors, ExNs and InNs, isoform usage varied most in cell subtype specification and developmental stages, with only a few isoform variations in disease conditions. In contrast, vascular and microglia cells showed more isoform variability distribution in disease-specific contexts, suggesting a potential role of isoform switch regulation in these cell types in the context of ASD (fig. S7B).

Given that neurodevelopmental diseases show the least variability in neurogenic cell types (Progenitor and ExN), we conducted further investigations into isoform regulation between ASD and WT. We calculated differential transcript usage (DTU) for each cell type using the R package IsoformAnalyzeR. As expected, CPNs harbored the most genes with DTUs (n = 255) and there were no DTU events detected in IPCs (Fig. 5A, table S5). This finding is consistent with our previous analyses indicating CPNs exhibit the most DEGs and DEIs between WT and ASD (Fig. 4J). While ASD-associated DEGs and DEIs of each cell type were significantly enriched in the corresponding cell type marker genes, DTUs (represent isoform switching) only significantly aligned with cell type markers in CPNs (Fig. 5B). This suggests that ASD-related alternative splicing is highly attuned to cell type-specific functions, especially in the upper-layer neurons.

**Fig. 5.**
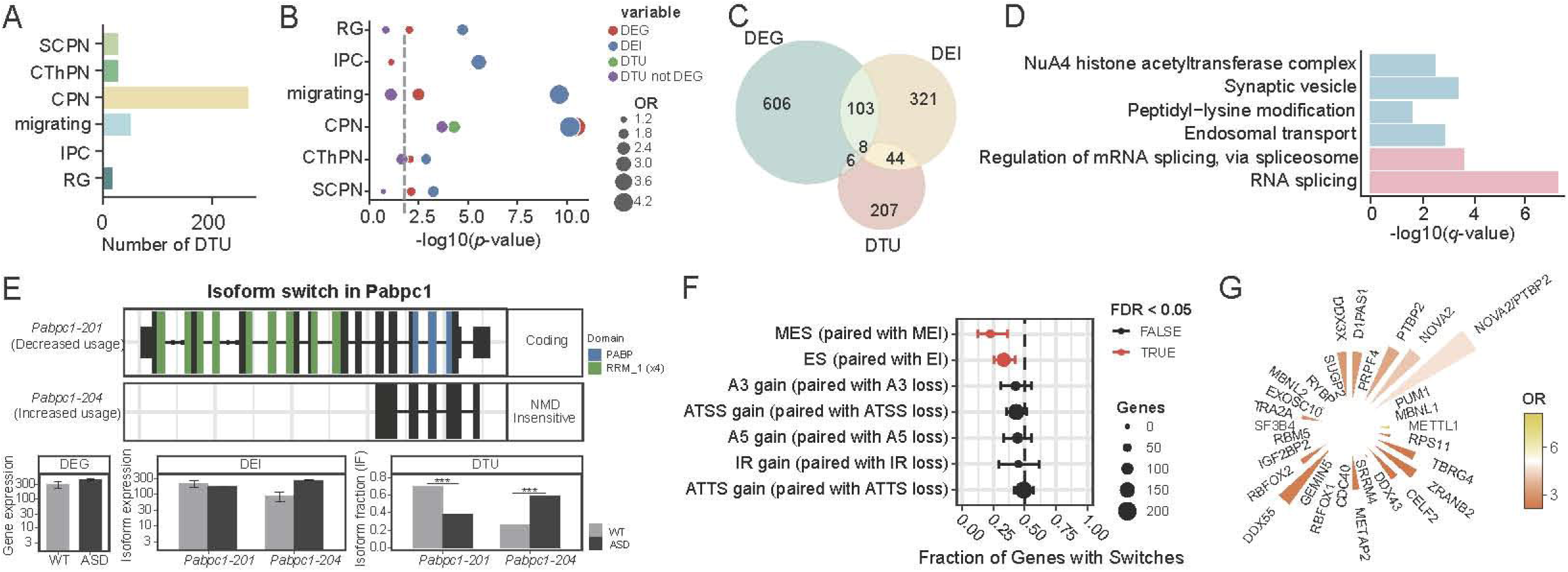
The landscape of isoform variability in development and disease. (**A**) Numbers of ASD related DTUs in each cell type. (**B**) Comparison of ASD features including DEGs, DEIs, DTUs and DTUs but not DEGs with corresponding cell type markers in each cell type. The x-axis represents the p-values obtained from Fisher’s exact test. The circle size denotes odds ratio (OR), reflecting the strength of the association between ASD features and cell type markers. (**C**) Overlap of ASD related DEG, DEI genes, and DTU genes in CPN. (**D**) GO terms of DTU genes in CPN. (**E**) Example showing isoform switch of *Pabpc1* gene in CPNs between WT and ASD. The top panel shows isoform structure of *Pabpc1-201* and *Pabpc1-204*. RABP: polyA-binding protein domain; RRM: RNA recognition motif; NMD: nonsense-mediated mRNA decay. The bottom panels present expression abundance and fraction of the two isoforms in WT and ASD CPNs. (**F**) Enrichment/depletion in isoform switch consequences.The fraction (and 95% confidence interval) of isoform switches (*x*-axis) resulting in gain of a specific alternative splice event (indicated by *y*-axis) in the switch from WT to ASD. Dashed line indicates no enrichment/depletion. Red indicates if FDR<0.05. Splicing abbreviation is as in Table S8. (**G**) Bar plot depicting enriched RBPs for the DTU genes in CPN. The height of the bar represents -log10(*p* value) of fisher exact test, and the colors represent odds ratios (OR) for enriched RBPs.

Further comparison of these DTU genes with DGE and DEI genes in CPNs Revealed minimal overlap, with only 6% (14/221) and 23% (952/221) DTU genes overlapping with DEGs and DEIs, respectively (Fig. 5C). The 207 DTU unique genes were significantly enriched in RNA splicing pathways, suggesting that CPNs in ASD might suffer from splicing dysregulation (Fig. 5D). This is in line with findings that altered RNA splicing can precipitate neurodevelopmental disorders (*20, 38, 57*). The isoform switching genes were also enriched in endosomal transport and synaptic vesicle function, processes critical for maintaining normal synaptic activity and neuronal communication. For example, *Pabpc1*, which stabilizes mRNAs by binding to the poly(A) tail and is essential for regulating protein synthesis in neurons (*80, 81*), exhibited a significant shift in isoform usage (Fig. 5E). The functional mRNA-binding isoform *Pabpc1-201*, containing RNA recognition motif (RRM) and a poly(A)-binding protein domain (PABP), was dominantly used in WT CPNs. Conversely, in ASD, the usage of this isoform was significantly reduced, with an increase in the usage of another isoform *Pabpc1-204*, which lacks coding potential and is insensitive to nonsense-mediated mRNA decay (NMD) (Fig. 5E). The switch from the functional *Pabpc1-201* to the non-functional *Pabpc1-204* isoform may disrupt mRNA regulation and translation, contributing to the developmental abnormalities observed in CPNs in ASD.

Considering the widespread alterations in isoform usage in ASD CPNs, we investigated the underlying mechanisms driving these changes. By analyzing the AS patterns of DTU isoforms in ASD CPNs, we found multi-exon inclusion (MEI) and exon inclusion (EI) significantly contributed to the upregulated isoforms in ASD (Fig. 5F). As RNA-binding proteins (RBPs) play a crucial role in controlling AS of RNAs (*2, 82*), we examined the enrichment of the DTU genes among different RBP targets collected from brain and cancer samples (*24*). A total of 29 RBPs were identified as potentially involved in the aberrant splicing of DTU genes in ASD (Fig. 5G, table S6). Among these, nine were identified as ASD risk genes, such as MBNL1 (*83*), DDX3X (*84*)and CELF2 (*85*), while eight were reported to be associated with neurological diseases (*86–88*). These splicing regulators may be central to unraveling the molecular mechanisms underlying ASD risk genes.

### Loss of isoform diversity in ASD related to epigenetic factors

We also investigated whether there were changes in isoform diversity along neurogenesis in ASD. Similar to the observations in WT samples, the isoform diversity was lowest in IPCs (Fig. S7C). However, a significant loss of isoform diversity was observed in ASD compared to WT (∼47% genes expressing multiple isoforms in ASD versus ∼55% in WT) (Fig. 6A). This reduction indicates a broad dysregulation of gene splicing across neurogenesis stages and suggests a potential impact on the functional complexity of the transcriptome. Strikingly, SCPNs and IPCs in ASD exhibited the most substantial loss of isoform diversity (Fig. S7D). Genes losing isoform diversity in ASD were enriched in biological processes such as autophagy, dendrite development, and DNA repair (Fig. S7E), with autophagy being the most significantly enriched GO term. The impairment of autophagy, as previously indicated, is a critical factor in neurodevelopmental disorders like ASD, potentially leading to the accumulation of damaged proteins or organelles and adversely affecting brain development and synaptic plasticity (*89–92*).

**Fig. 6.**
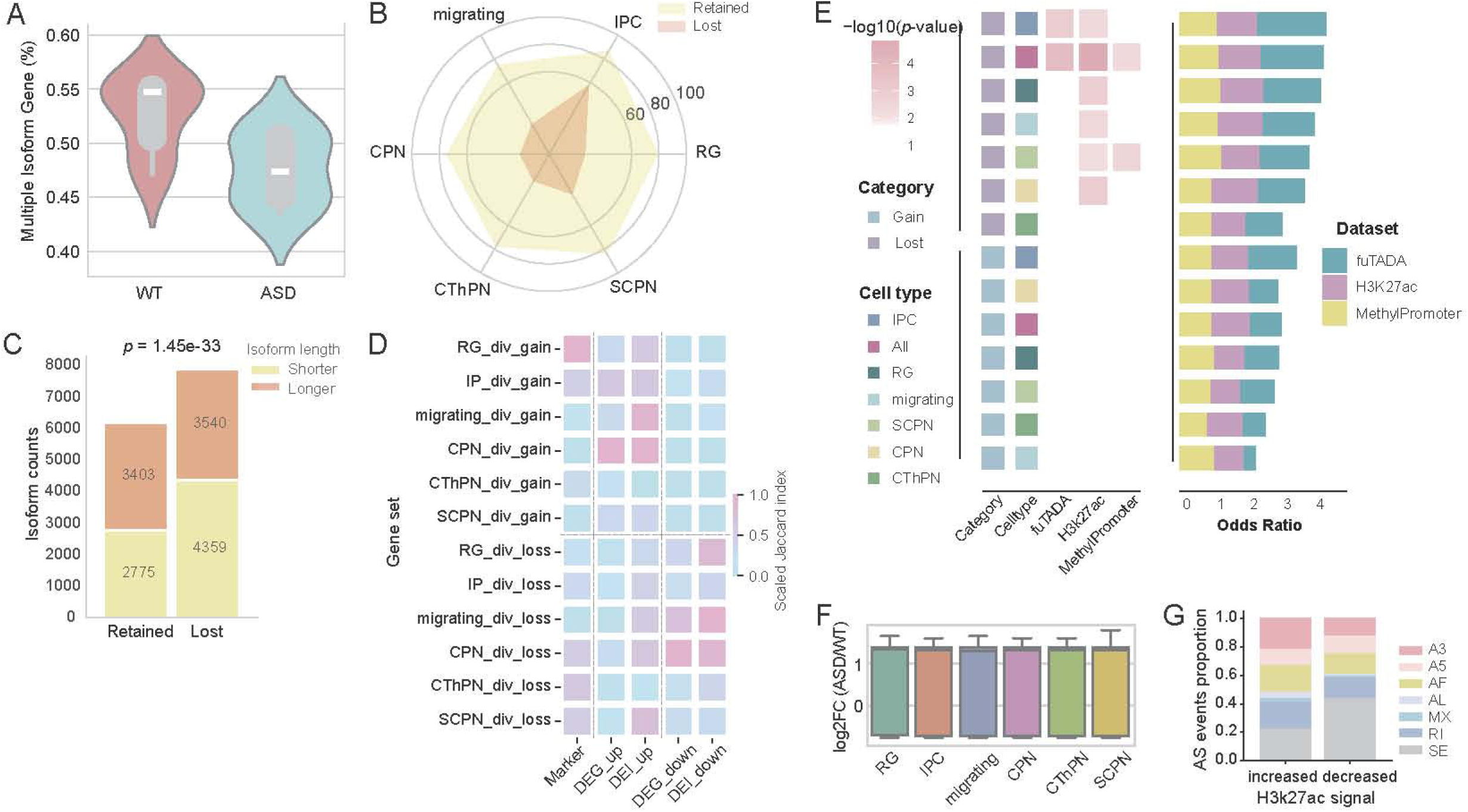
Loss of isoform diversity in ASD. **(A)** Violin plot showing the percentage of genes expressing multiple isoforms in WT and ASD. **(B)** Radar plot displaying the percentage of high-expression isoforms(log (TPM+1) > 10) in either retained or lost isoforms of ASD across different cell types. **(C)** Bar plot showing the number of isoforms categorized by length (relative to the median length of all isoforms) in retained versus lost isoforms in ASD. **(D)** Heatmap of the scaled Jaccard Index illustrating the overlap of genes gaining or losing isoform diversity in ASD and cell type markers, DEGs and DEIs between WT and ASD in each cell type. **(E)** Heatmap showing the enrichment significance of genes showed isoform diversity changes compared to three different types of ASD datasets across different cell types and overall. fuTADA contains ASD Rare Variants; H3K27ac, ASD-abnormal H3K27ac region; MethyPromoter: Promoter regions with altered DNA methylation levels in ASD. The collective odds ratio relative to the three datasets are shown on the right, highlighting the enrichment of each gene set in ASD-associated variants. **(F)** Boxplot depicting the changes of H3K27ac modification levels in genes with isoform diversity loss in ASD. **(G)** Bar plot illustrating the ratio of each AS event identified from isoform diversity-loss genes showing ASD-increased and ASD-decreased H3K27ac levels respectively. ES, Exon skip; RI, Retained intron; MX, Mutually exclusive exons; AL, Alternative last exon; AF ,Alternative first exon; A5, Alternative 5’ splice site; A3, Alternative 3’ splice site.

To further characterize the lost isoforms in ASD, we compared them with the retained isoforms within the same gene across cell types. The lost isoforms generally displayed low abundance in WT (Fig. 6B, fig. S7F) and were significantly shorter than the retained isoforms of the same genes (Fig. 6C, fig. S7G). These characteristics suggest that while the lost isoforms may contribute minimally to overall gene function under normal conditions, their absence in ASD could disrupt the regulatory balance or affect specific cellular processes by reducing isoform diversity and transcriptional noise.

We then examined the overlap of genes that experienced gained and lost isoform diversity in ASD with ASD-DEGs and ASD-DEIs, respectively (Fig. 6D). Genes losing isoform diversity closely resemble the DEGs and DEIs in migrating neurons and CPNs (Fig.6D). These results suggested that, despite CPNs showing fewer diversity-loss genes than SCPNs (fig. S7D), these genes in CPNs underwent more pronounced expression changes.

To investigate whether the changes of isoform diversity related to genetic or epigenetic alterations in ASD, we integrated data from fuTADA (*93*) and previously published data on ASD-associated abnormal H3K27ac (*94*) and promoter DNA methylation (*95*) (Fig. 6E). The genes losing isoform diversity were significantly enriched in regions with abnormal H3K27ac in ASD across multiple cell types, suggesting the splicing dysregulation might be influenced by enhancer activities across different neural cell types. Further comparison of H3K27ac modification levels in genes losing isoform diversity revealed a widespread elevation in H3K27ac level (Fig. 6F). We then separated genes lossing isoform diversity by the changes of H3K27ac level in ASD, obtaining 338 genes with increased H3K27ac level and 226 genes with decreased H3K27ac level. We then evaluated how different types of alternative splicing contribute to these genes in ASD. Interestingly, we noticed that the lost isoforms from genes with increased H3K27ac levels predominantly consist of exon skipping (44%) (Fig. 6G). These findings suggest that enhancer activity, as marked by H3K27ac, may be responsible for the loss of isoform diversity in ASD and these epigenetic disruptions may play a pivotal role in the pathophysiology of ASD.

## DISCUSSION

In this study, we employed a combination of TGS-based single-cell full-length isoform sequencing and newly transcribed RNA labeling to explore the isoform expression landscape in the developing mouse cortices at various embryonic stages. Our findings provide novel insights into cell-type-specific isoform diversity, isoform switching events, and the transcriptional dynamics that underpin cerebral cortex development. Leveraging new RNA expression data, we discovered that cell fate determination occurs earlier than inferred from total RNA profiles. IPCs, based on nascent RNA expression, already exhibit transcriptomic signatures indicative of migrating neurons, suggesting a primed state for differentiation. This underscores the utility of nascent RNA analysis in capturing early transcriptional events that shape lineage trajectories, affording a more precise temporal resolution of developmental processes. In addition, we identified previously unreported isoforms that were masked by total RNA analysis, which warrants further investigation in future studies. Furthermore, by examining the autism-associated isoform changes in the BTBR mouse model, we revealed dysregulated splicing and reduced isoform diversity in ASD. These findings provide a comprehensive view of the transcriptomic complexity in cortical development and its disruption in neurodevelopmental disorders.

Our analysis revealed that isoform diversity dynamically shifts during neuronal development, particularly in IPCs, which exhibited the lowest isoform diversity. This transient reduction likely reflects streamlined transcriptional activity associated with cell cycle progression, as supported by enrichment of cell cycle-related genes. In contrast, isoform diversity increased during neuronal differentiation, aligning with the activation of genes involved in neuron projection morphogenesis and synaptic function. The persistence of multi-isoform expression in certain genes underscores their critical roles across developmental stages, providing a nuanced view of isoform usage in neural specification. Our study also demonstrated the critical role of alternative splicing and isoform switching in the precise regulation of neurogenesis. We observed dynamic changes in isoform expression across multiple cell types and developmental stages, with progenitor cells and excitatory neurons showing the highest isoform diversity. These findings are consistent with previous reports indicating that splicing regulation is essential for neuronal differentiation and maturation (*19, 96*). Specifically, we identified isoform switching in genes such as *Ergic3* and *Clta*, which are known to regulate intracellular transport and synaptic plasticity, respectively. These events may fine-tune the functions of proteins in a cell-type-specific manner, allowing for specialized roles during different stages of neurodevelopment. These findings contribute to a deeper understanding of how isoform diversity is maintained and modulated throughout neurogenesis, complementing existing research on splicing regulation in brain development (*1, 97*).

Our data suggest that isoform diversity is crucial for maintaining the functional complexity of the developing cortex (*24*). Loss of isoform diversity in specific cell types, such as CPNs, may impair the brain’s ability to adapt to functional demands during development (*98*). ASD cortices exhibited a notable reduction in isoform diversity and the corresponding genes were enriched in processes such as autophagy, dendrite development, and DNA repair, which are critical for maintaining neuronal integrity. Although the lost isoforms were generally shorter and less abundant, their loss may impair neural complexity, contributing to the pathological features of ASD. By further integration with epigenetic data in ASD, we found a correlation between reduced H3K27ac signal and increased exon skipping events, indicating the dysregulation in RNA splicing be associated with aberrant H3K27ac levels. This is consistent with recent research highlighting the role of epigenetic modifications in regulating alternative splicing during brain development (*99–101*). But how changes in chromatin state influence transcription splicing needs further exploration.

We also observed an aberrant temporal shift in CPN generation in ASD embryonic cortices, with premature emergence at E13.5 compared to E15.5 in WT cortices. This premature differentiation was coupled with a less mature pseudotime score, suggesting disrupted neurogenesis timing. These findings support the hypothesis that early dysregulation in neuronal lineage specification contributes to cortical abnormalities observed in ASD. The key genes in the progenitors influencing the abnormal proportions of CPNs include mitochondrial function-related genes (*Atp5g1*, *Uqcrh*), implicating energy metabolism in these processes. Isoform-level expression analyses revealed widespread dysregulation across cell types, with CPNs exhibiting the most ASD-associated DEIs. Dysregulated isoforms were enriched in pathways associated with synaptic function, mitochondrial activity, and mTORC1 signaling, aligning with known ASD-related mechanisms. The identification of specific isoforms, such as *Tcf4* and *Pabpc1*, highlights how isoform switching alters critical neuronal processes, such as transcriptional regulation and mRNA stabilization, potentially contributing to ASD pathology. RBPs were implicated as key regulators of these splicing abnormalities in ASD CPNs, with several identified as ASD risk genes, including *MBNL1*, *DDX3X*, and *CELF2*. The role of RBPs in ASD highlights a mechanistic link between splicing regulation and neurodevelopmental disorders. Targeting RBPs or correcting specific splicing events may offer therapeutic potential for ASD.

Pseudogenes have gained increasing attention in recent years due to their potential regulatory roles in various biological processes and diseases, including ASD (*20, 102*). The advent of TGS-based full-length isoform sequencing has provided unprecedented opportunities for investigating pseudogenes by enabling direct identification of full-length transcripts, even in highly similar genomic regions. This technology excels in distinguishing pseudogenes from their functional counterparts, detecting low-abundance transcripts, and uncovering alternative splicing events. Notably, we identified significant pseudogene dysregulation, with 12.9% of common DEIs attributed to pseudogenes. These noncoding transcripts may act as competitive endogenous RNAs, modulating the expression of their parent genes. The complex interactions between pseudogenes and parent genes warrant further investigation to elucidate their roles in ASD pathology.

While our study provides a comprehensive overview of isoform expression during *in vivo* neural development in WT and ASD cerebral cortices, this study shows some limitations. Most importantly, the relatively shallow sequencing coverage of TGS may have limited the detection of lowly expressed isoforms, which could provide additional insights into splicing regulation. Future studies using higher sequencing depth or complementary methods such as targeted isoform enrich may help overcome this limitation. In conclusion, we have provided a high-resolution view of isoform expression dynamics during cortical development and our study underscores the importance of isoform diversity in cortical neurogenesis and its disruption in early ASD development. The observed reduction in isoform diversity and splicing dysregulation in ASD points to alternative splicing as a critical mechanism underlying the molecular pathology of the disorder. Future research aimed at restoring splicing diversity in ASD may hold therapeutic potential for alleviating the neurodevelopmental deficits associated with the condition.

## MATERIALS AND METHODS

### Mice and in utero surgeries

All animal experiments were approved by the Institutional Animal Care and Use Committee, Guangzhou laboratory (Guangzhou, China). C57BL/6 mice were obtained from Baishitong company (Zhuhai, China) and mated overnight. BTBR T+ Itpr3tf (BTBR) mouse were obtained from The Jackson Laboratory (the United State). The morning that plugs were detected was considered E 0.5. All mice were housed in specific pathogen-free facilities, maintained on a 12 h light/dark cycle with controlled temperature and air humidity, and allowed free access to food and water. The research license number is GZLAB-AUCP-2022-08-A01.

Timed pregnant mice were anaesthetized by intraperitoneal injection with 0.015 ml/g body weight of 2.5% tribromomethyl alcohol and 2.5% tert-amyl alcohol in phosphate buffered saline (PBS). In utero surgery and injection of 4-thiouridine (4sU, 500mM, sigma) in the lateral ventricles of the embryonic mouse forebrains at E13.5-E17.5 were performed. A capillary glass needle was used to inject approximately 1.5 µl of the 4sU per embryo.

### Tissue preparation

The pregnant mice treated with 4sU were sacrificed after four hours by cervical dislocation. Then embryonic mouse prefrontal cortex were dissected. For single-cell suspension preparation, dissected embryonic mouse prefrontal cortex were homogenized and lysed in 500ul HBSS buffer containing 30U/ml papain and 2ul DNase. The tissue was incubated at 30℃ for 5-10 min and gently pipetted, then was terminated digestion with 500ul cooled hibernate E. The solution was filtered using a 40-μm cell strainer (Falcon cat. no. 352340) and collected in a tube, and centrifuged at 300g for 3 min at 4 °C. The supernatant was removed and the pellet resuspended in 0.2 ml PBS with 0.01% BSA. The cell lysis was analysed with cell viability and cell number. Only cell viability more than 90% was used for the next process.

### Single-cell transcriptome amplification and NGS library construction for Illumina sequencing

For labeling experiments, according to the protocol of the DynaSCOPE® Single Cell Dynamic RNA Library Kits (Singleron), single-cell suspensions (2×10^5^ cells/mL) with PBS (HyClone) were loaded onto microwell chip using the Singleron Matrix® Single Cell Processing System. Barcoding Beads are subsequently collected from the microwell chip, followed by reverse transcription of the mRNA captured by the Barcoding Beads to obtain cDNA, and PCR amplification. The amplified cDNA is then fragmented and ligated with sequencing adapters. The scRNA-seq libraries were constructed according to the protocol of the GEXSCOPE® Single Cell RNA Library Kits (Singleron). Individual libraries were diluted to 4 nM, pooled, and sequenced on Illumina NovaSeq system with 150 bp paired end reads.

### TGS library construction for PacBio sequencing

We used the same amplified cDNA from above for each embryonic sample. Briefly, eighty nanograms of cDNA products were amplified for another five PCR cycles using KAPA HiFi HotStart Uracil 2 x ReadyMix (Kapa Biosystems, Cat. KK2602) and designed primers Biotin-ACTAG/ideoxyU/CTACACGACGCTCTTCCGATCT and ACTAG/ideoxyU/AAGCAGTGGTATCAACGCAGAG. The PCR products were then purified using 0.8 volumes of Agencourt AMPure XP Beads (Becman, Cat. A63882), quantified using Qubit dsDNA HS Assay Kits (Thermo Fisher, Cat. Q32854). The barcode-UMI-poly (dT)-flanked cDNAs were captured on streptavidin-coated M-280 Dynabeads using Dynabeads^TM^ kilobaseBINDER^TM^ Kits (Invitrogen, Cat. 60101,). After washing, the beads were re-suspended in a 19 μL reaction buffer containing 2 μL 10× T4 DNA ligase buffer (NEB, Cat. B0202S) and 1 μL USER Enzyme (NEB, Cat. M5505) to nick, then 1ul T4 DNA ligase (NEB, Cat. M0569) to ligate inserts. The resultant multi-insert library was purified using 0.4 volume of Agencourt AMPure XP Beads and was then end-repaired and A-tailed using the NEBNext Ultra II End Repair/dA-Tailing Module, with incubation for 15 min at 20 °C and then for 30 min at 65 °C. The cDNA was ligated with 2 μL of a pre-annealed 10uM dT-overhang selection adapter (GAACGACATGGCTACGATCCGACTT; Pho-AGTCGGATCGTAGCCATGTCGTTC) using the NEBNext® Ultra™ II Ligation Module. Then, 100 ng of the purified products was PCR-amplified for 8-9 cycles using KAPA HiFi HotStart 2x ReadyMix (Kapa Biosystems, Cat. KK2602). The amplified products were again purified using 0.4 volume of Agencourt AMPure XP Beads to produce the SMRTbell Template. To remove residual adapters and unligated DNA fragments, exonuclease and NEBuffer 3 (NEB) were added to the library before incubation at 37 °C for 1 h. The products were purified using 0.8 volume of Agencourt AMPure XP beads, eluted with 15 μL elution buffer (10 mM Tris-HCl, pH 8.0), and quantified. Sequencing primer annealing and polymerase binding to the PacBio SMRTbell Templates were performed according to the manufacturer’s recommendations (PacBio, US). The library complex was then sequenced using SMRT Cell 8M on sequel IIe system (PacBio).

### RT-QPCR

Total RNA was extracted from cerebral cortices tissues using RNeasy Mini Kit (Qiagen) according to the manufacturer’s instructions. The RNA was quantified by absorbance at 260 nm using Nanodrop (Thermo Scientific). cDNA was synthesized by HiScript III RT SuperMix for qPCR (+gDNA wiper) (Vazyme, R323) from total RNA. QPCR was carried out using Quantagene q225 (Kubo) in triplicate. The expression data were normalized to glyceraldehyde-3-phosphate dehydrogenase (*Gapdh*) mRNA expression.

### Immunofluorescence staining

The pregnant mice were sacrificed by cervical dislocation and embryonic mouse brains were dissected. For paraffin sections, tissue was postfixed for 24 h in 4% PFA at 4°C, then processed and embedded in paraffin, and cut into 5 mm sections for immunofluorescence analysis, which was performed as previously described. In brief, coronal sections (5mm) encompassing the prefrontal cortex were serially collected. Sections were pre-incubated in a blocking solution containing 5% normal donkey serum and 0.3% Triton X-100 in 50 mM TBS, pH 7.4, at room temperature for 1 h. Primary antibodies were diluted in 1% BSA and 0.3% Triton X-100 in TBS, pH 7.4, and incubated overnight at 4°C. The following primary antibodies were used: . After three washes, the sections were then incubated with the secondary antibodies (Thermo Fisher Scientific), which were conjugated with Alexa-488, Alexa-555, or Alexa-647, at room temperature for 1 h. Finally, sections were visualized under a confocal laser scanning microscope (LSM 980, Carl Zeiss).

### Quality control and clustering of the scRNA-seq data

Short-read scRNA-seq data were imported into Python package Scanpy (version 1.9.3), genes and cells were filtered by the criteria: genes with expressed cells > 3, unique genes > 1000, 2000<counts<40000 and percent mitochondrial genes < 20. After filtering, we received 55773 cells with NGS gene expression files. We then normalized the gene measurements for each cell by the total mapped reads, multiplying this by a scale factor (10000) and log+1-transformation of the result. We then selected the top 2000 most variable genes with the selection method ‘vst’ for downstream PCA. The first 30 latent representation was then used for dimensionality reduction via UMAP and Louvain clustering. Resolution for Louvain clustering was set at 1.5 in order to obtain fine clusters. To annotate clusters, we determined DEGs using rank_genes_group function (Wilcoxon rank-sum method) for each cell cluster. We required genes to express in over 25% of the cells within the cluster and at a minimum of 0.25-log fold change. By reviewing the resulting markers as well as the expression of canonical marker genes, we assigned the cell-type identity to every cluster.

For TGS data, we generally followed the preprocessing process described above. However, due to lower depth of long-read sequencing, we used more loose quality control standards (unique genes > 500, 1000<counts<40000). We then kept cells that were captured in both NGS and TGS datasets, therefore, the cell type annotation would be transferred to TGS data based on cell barcodes. For all WT samples, we received 12418 cells for isoform expression analysis. We chose 30 PCs of isoform-expression data for downstream clustering analysis.

### Preprocessing of PacBio HiFi sequencing data

To process Pacbio data in our study, we followed the pipeline introduced by scISA-Tools, which can be found at https://github.com/shizhuoxing/scISA-Tools. Briefly, to obtain circular consensus sequencing (CCS) reads, we utilized SMRT-Link (v10.2.1.143962) and applied the following parameters: “--min-passes 1 --min-length 50 --max-length 21000 --min-rq 0.75”. To identify full length non-chimeric (FLNC) sequences from the CCS reads, we used NCBI BLAST(v2.10.0) to locate 5’ and 3’ primers with the following parameters:”-outfmt 7 -word_size 5”. Specifically, we generated FLNC reads with a minimum primer alignment length of 16 bp and a minimum transcript length of 50 bp. Next, we performed an alignment of FLNC reads to the mouse genome CRCm39 using minimap2(version 2.17-r941) employing the splice-aware alignment mode specifically designed for PacBio Iso-Seq data with parameters “-ax splice -uf --secondary=no -C5”. Following the alignment, we employed gffcompare (v0.11.6) to filter and retain only the uniquely mapped exonic reads based on GENECODE vM33 with a class code of "=ckmnjeo". To enhance the accuracy of our analysis, we incorporated corrected cell barcodes and unique molecular identifiers (UMIs) with a strategy similar to 10× Genomics CellRanger. Gene-level expression matrix was finally generated. For isoform quantification, we used Isoquant (version 3.2.0) with parameters ‘--data_type pacbio_ccs’ to precisely map reads to reference transcripts. We kept reads mapped to only one certain transcript.

After obtaining the gene and isoform expression level, we identified newly synthesized mRNA captured by 4sU-labeling experiments. Generally, we identify the T-to-C (A-to-G for reverse strand) transition in every read by comparing with the reference genome. When processing every labeled sample, we subtracted the T-to-C SNP loci information generated from non-labeled scRNA-seq of the same sample in order to elevate accuracy. If reads with the same UMIs show at least one substitution, the transcripts indexed by the UMIs were regarded as new RNA molecules. The new and old transcripts were counted, then we could obtain digital matrices with new and old quantification at both gene and isoform levels.

### Preprocessing of short-read 4sU labeled scRNA-seq

We utilized dynaseq function developed in CeleScope (v1.6.1) to generate short-read transcriptomics matrix as well as to identify newly synthesized mRNA by calling T-to-C SNPs compared to mouse genome CRCm39. We used default parameters for this analysis.

### RNA velocity analysis for 4sU-labeled data

We used Dynamo package for metabolic RNA-velocity analysis on E17.5-1 sample only which contains 3 replicated embryo samples, at isoform resolution (*15*). Six main neurogenic cell types were kept for further analysis. The UMAP was created by parameters: top genes equaling to 4000, pca components to be 30, and nearest neighbors to be 50 with preprocessing recipe ‘Seurat’. With one-shot 4sU label time to be 4, the later velocity analysis was forced from neurogenesis start point RG to two terminal states, which are CPNs and CFuPNs, with function ‘confident_cell_velocities’. The two Least action paths (LAP) were found between fixed points, which indicate the most possible transition probability from progenitor to two target cell types. Along each LAP, isoforms of mouse TFs with largest with top 1000 mean square displacement (MSD) were ordered by their maximum expression occurrence time.

### Total and new RNA expression comparison

New and total gene expression matrices were merged for dimension reduction analysis followed by standard single-cell processing procedures with 30 pca components and 50 nearest neighbors. Since the nascent RNA compromise ∼20% of the total RNA expression according to NGS 4sU efficiency, we downsampled the NGS data to 20% and did similar analysis with the original new matrix. For isoform matrices, we applied trajectory analysis with merged data using pyVIA package which shows composition of clustered cells along trajectory.

In order to find the reason why the two lineages split at IPC, we used similarity comparison methods to confirm that largest differences exist in the two matrices in IPCs and migrating neurons. Specifically, six machine learning classifiers (RF, MLP, SVM, DT, GB, LR) were trained with total RNA expression profiles. The input data was total isoform expression of single cells while the labels were the four cell type categories which are RG, IPC, migrating and ExN. After training, we tested the classifiers by new RNA expression of single cells. Tested targets were compared with true labels to calculate confusion matrix. We then concatenate scaled new and total expression of each isoform into arrays and grouped trends with hierarchical clustering. With cosine metric and complete linkage, we obtained 30 dynamic clusters of expression.

### ASD associated DEG and DEI analysis

Between ASD and WT samples, we utilized pseudo-bulk strategies for every cell type. We used DESeq2 to perform DEI analysis. We kept isoforms with p-adjust values smaller than 0.05 and log2 fold changes larger than 0.5. Similarly, we regarded genes with p-adjust values smaller than 0.01 and log2 fold changes larger than 1 as DEGs.

### Correlation analysis to identify genes related to CPN proportion

We referred to the work of Jourdon, A. et al. (*45*) to correlate the gene expression of three upstream progenitor cell types (RG, IPC, and migrating cells) with the CPN proportion in WT and ASD libraries (n=7). For gene expression, we used the average log10 (CPM)-normalized expression values per sample within each progenitor cell type. For each pair of gene and CPN proportion across progenitor cell types, we computed Spearman’s correlation coefficient and corresponding p-value. We extracted candidate genes with *p*-values < 0.05 and positive correlation coefficients.

### TF enrichment analysis

Possible TFs that regulate DEIs were enriched by the enrichR package based on TRANSFAC_and_JASPAR_PWMs, ChEA_2022 and ENCODE_TF_ChIP-seq_2015 databases. ASD associated DEIs were split into two groups by up-or down-regulation. If the enrichment *p*-value is smaller than 0.05 and the TFs were expressed in corresponding cell types, they were kept. For visualization, we only kept TF isoforms with top 100 odds ratio.

### Transcript usage and variability analysis

For marker isoforms in WT samples, we found two patterns of isoform switch scenarios in DEI. We extracted isoforms when they showed different directions of regulation in Progenitor and EX. Then we classified the isoform switch event as classical if the usage of the dominant isoforms flip compared with the other isoforms of the same gene. Otherwise if the isoform usages ratio did not reverse, this event was defined as a non-classical switch event.

We adapted the method of Joglekar, A. et al. (*11*) to assess isoform usage variability along three axes: development (embryonic stages), cell type (specific subtypes of the five main cell types shown in Fig. 2), and disease state (WT or ASD). To calculate variability, we first determined the percent inclusion (PI) for each isoform along each axis. For a given axis (e.g., development), the PI was computed by summing inclusion values across relevant conditions within that axis (such as all cell types for development variability). The variability for each isoform was then defined as the difference between the maximum and minimum PI values along that axis. This provided a measure of how much isoform usage fluctuated across stages, cell types, or disease states. An isoform was excluded from further analysis if it lacked a PI value for any axis. Isoforms exhibiting at least a 10% variability (PI ≥ 0.1) across any of the three axes were retained for further analysis. The values were normalized to sum to 1 and visualized in a ternary plot. If the normalized disease state variability exceeded 0.5, while developmental and cell type variability remained below 0.5, the isoform was considered disease-specific for a particular cell type.

We used the R package ‘IsoformAnalyzeR to calculate’ DTU of every cell type between ASD and WT. Isoform switch events were identified if the usage changes were larger than 0.1. We summarized the alternative splicing events with analyzeSwitchConsequences function.

### Isoform diversity analysis

We kept isoforms which are expressed in over 10 cells in WT and 3 cells in ASD. For every gene, we summarized the number of isoforms that are expressed. When the expressed isoforms are larger than 2, the diversity category was set to be ‘multiple’; if a single isoform was expressed, the diversity category was set to be ‘single’; else, the gene was ‘not expressed’.

To compare the expression levels of ASD-retained and ASD-lost isoforms, we first converted the expression of all isoforms to averaged log (TPM+1) in WT pseudo-bulk samples. We sorted the expression values of each isoform in a descending order and removed the top 10% isoforms to minimize the impact of highly expressed outliers. Then expression values were divided into 10 bins and isoforms with log (TPM+1) < 10 were classified as low-expressed, while those with values exceeding this threshold were considered high-expressed. For each cell type, we calculated the proportion of high-expressed isoforms within both retained and lost categories by dividing the count of high-expressed isoforms by the total isoforms in each category. This result was plotted at Fig6B and FigS7D.

### Isoform length analysis

To assess the length differences between ASD-retained and ASD-lost isoforms within the same genes, we only extracted genes expressing multiple isoforms in WT and single isoforms in ASD. The length of each isoform was referred to the transcript annotation from the Gencode vM33 annotation. Then we calculated the mean lengths of the isoforms belonging to the two groups within each gene and then used a paired t-test to evaluate the statistical significance between the mean lengths of the two groups of isoforms (FigS7G).

Meanwhile, a Chi-Square Test was performed to compare the length difference between ASD-retained and ASD-lost isoforms within the same genes. All isoforms were ranked according to the length and isoforms were categorized as ‘longer’ if their length exceeded the median length of all isoforms, otherwise they were categorized as “shorter”. Then we counted the numbers of longer and shorter isoforms in the two groups of isoforms and the chi-square test was performed (Fig6C).

### Isoform enrichment analysis

Genes showing changes in isoform diversity were classified into two categories: "diversity gain" and "diversity loss." To evaluate the expression effects of these changes, we used the Jaccard similarity index to assess the overlap between these genes and ASD-DEGs, ASD-DEIs, and cell type markers respectively.

To investigate potential epigenetic factors influencing isoform diversity changes, we used Fisher’s exact test to assess the overlap between the diversity-gain and diversity-loss genes and those associated with genetic and epigenetic variations. ASD rare variant genes were obtained from the fuTADA database (*93*), while epigenetic factors were assessed using published data including H3K27ac changes (*94*) and promoter methylation changes (*95*) in human ASD samples compared to WT samples. Thresholds of *p* < 0.05 were used to define significant enrichment.

For genes that showed isoform diversity loss as well as aberrant H3k27ac modification in ASD, we extracted the fold changes of H3K27ac levels in ASD samples compared with WT samples. After that, we divided these genes into two groups based on up-regulated or down-regulated H3K27ac modification. Then the AS events were analyzed by comparing the ASD-lost isoforms with ASD-retained isoforms within each gene. The AS events reference was the mouse genome (ioe format) generated from SUPPA2.

